# GdDO3NI allows imaging of hypoxia after brain injury

**DOI:** 10.1101/2021.03.16.435723

**Authors:** Babak Moghadas, Vimala N. Bharadwaj, John P. Tobey, Yanqing Tian, Sarah E. Stabenfeldt, Vikram D. Kodibagkar

## Abstract

**Purpose:** In this study, we use the hypoxia targeting agent (GdDO3NI, a nitroimidazole-based T_1_ MRI contrast agent) for imaging hypoxia in the injured brain after experimental traumatic brain injury (TBI) using magnetic resonance imaging (MRI), and validate the results with immunohistochemistry (IHC) using pimonidazole.

**Methods:** TBI induced mice (controlled cortical impact model) were imaged at 7T using a T_2_ weighted fast spin-echo sequence to estimate the extent of the injury. The mice were then were intravenously injected with either conventional T_1_ agent (gadoteridol) or GdDO3NI at 0.3 mmol/kg dose (n=5 for each cohort) along with pimonidazole (60 mg/kg). Mice were imaged pre- and post-contrast using a T_1_-weighted spin-echo sequence for three hours. Regions of interests were drawn on the brain injury region, the contralateral brain as well as on the cheek muscle region for comparison of contrast kinetics. Brains were harvested immediately post imaging for immunohistochemical analysis.

**Results:** GdDO3NI is retained in the injury region for up to 3 hours post-injection (p< 0.05 compared to gadoteridol) while it rapidly clears out of the muscle region. On the other hand, conventional MRI contrast agent gadoteridol clears out of both the injury region and muscle rapidly, although with a relatively more delayed wash out in the injury region. Minimal contrast enhancement was seen for both agents in the contralateral hemisphere. Pimonidazole staining confirms the presence of hypoxia in both gadoteridol and GdDO3NI cohorts, and the later cohort shows good agreement with MRI contrast enhancement.

**Conclusion:** GdDO3NI was successfully shown to visualize hypoxia in the brain post-TBI using T1-wt MRI.

## 1. Introduction

Traumatic brain injury (TBI) occurs due to damage to the brain resulting from an external mechanical force, including rapid acceleration or deceleration, blast waves, crushing force, an impact, or penetration by a projectile(1). An estimated 1.7 million TBI occur annually in the U.S., which results in the hospitalization of 373,000, and 99,000 totally disabled individuals, and over 50,000 deaths (2,3). Disabilities as well as cognitive, behavioral, and productivity problems are the most common complications for survivors; in acute cases, the affected individuals face mortality(4,5). Upon sustaining a TBI, the mechanical forces from impact inflict heterogeneous tissue damage, referred to as the primary injury phase(1). This insult initiates a myriad of pathophysiological and biochemical secondary injury signaling cascades, including hypo- or hyper-perfusion, edema, blood-brain barrier (BBB) dysfunction, and inflammation that evolves from minutes to days post-trauma(6–8).

Brain tissue hypoxia, defined as brain tissue oxygen tension (pO_2_) < 15 mm Hg, is a common consequence of TBI due to the rupture of blood vessels during impact(9). Studies have reported about 30-50% of TBI patients to have hypoxia as early as arriving in an emergency room(10,11). It is shown that brain hypoxia, as one of the post-traumatic insults, is associated with mortality and poor and unfavorable neurological outcomes(12–17). Tissue hypoxia has a significant cross-talk with inflammatory processes, whereby hypoxia can trigger the upregulation of proinflammatory cytokines, and the resulting inflammation can further exacerbate hypoxia due to an increase in the metabolic demands of cells and a reduction in metabolic substrates caused by thrombosis, trauma, compression (interstitial hypertension)(18). Cellular signaling continues acutely and post-acutely to restore homeostasis in injured tissue, which, if not controlled, can exacerbate the injury(8,19). Furthermore, studies suggest brain tissue oxygen-based therapy can reduce mortality rate and ameliorate neurological outcome for the patients (20,21).

Noninvasive characterization of the injury microenvironment is often difficult to achieve through conventional neuroimaging methods (e.g. Computed tomography (CT), T1w and T2w Magnetic Resonance Imaging (MRI)) as they are not sensitive enough to identify regions undergoing microstructural changes (22–24). The CT scan is highly effective in detecting TBI induced skull fractures, bleeding within and surrounding the brain (hematomas) as well as brain swelling (edema) and the resolution of these over time. CT is much more limited in its ability to detect the widespread microscopic injury to axons which leads to many of the long-term problems experienced by TBI patients. MRI is a powerful diagnostic tool that can detect signs of injury such as minute bleeding (microhemorrhage), small areas of bruising (contusion) or scarring (gliosis), which are invisible to the CT scan. Several MR-based neuroimaging modalities have been used to qualitatively examine acute and chronic changes post TBI longitudinally (25,26): (a) fluid-attenuated inversion recovery (FLAIR) MRI, a sequence that suppresses the high signal from CSF, is sensitive in detecting traumatic lesions and hematomas, (b)T_1_-weighted structural MRI is sensitive to morphological changes in gray matter volume and cortical thickness, (c) diffusion-weighted MRI (DWI) is sensitive to changes in the microstructural integrity of white matter, (d) MR spectroscopy provides a sensitive assessment of metabolic and neurochemical alterations in the brain, and (e) T_2_*-weighted blood oxygen level dependent (BOLD) functional MRI (fMRI) provides insight into the functional changes that occur as a result of structural damage and typical developmental processes. BOLD fMRI only reflects vascular oxygenation and it cannot provide data in regions where BBB is disrupted. An unmet need in TBI diagnosis is the ability to assess tissue oxygenation or hypoxia.

Among various invasive techniques to detect hypoxia, the 2-nitroimidazole pimonidazole, has been previously demonstrated to be reliable in detecting hypoxic regions in tissue (27). After intravenous administration and extravasation, pimonidazole gets activated under hypoxia and binds to thiol-containing proteins in hypoxic regions, and these adducts can be detected *ex vivo* utilizing immunohistochemistry (IHC) in conjunction with fluorescent microscopy (28). In addition, several noninvasive imaging approaches to assess hypoxia (qualitatively or quantitatively) have been developed and are in various stages of validation from preclinical to clinical use (29). Currently, [^18^F]Fluoromisonidazole (^18^F-MISO) [^18^F]fluoroazomycin arabinoside (FAZA) (30), [^18^F]-EF5 (31), [^18^F] fluoroerythronitroimidazole (FETNIM) (32) and Cu-labelled diacetyl-bis(*N*(4)-methylthiosemicarbazone (Cu-ATSM) are being used as hypoxia targeting PET imaging agents (33). While PET based probes have significantly advanced the field of hypoxia imaging, there is a strong rationale for the development of MRI based hypoxia-imaging techniques as well due to the ability of MRI to acquire higher resolution anatomical and complementary functional information in the same scanning session. BOLD and tissue oxygen level-dependent (TOLD) signal (34–37) and [^19^F] Tri-fluoromisonidazole (TF-MISO)(38,39) are MRI techniques to qualitatively image hypoxia while ^19^F (29,40) and ^1^H(41-43) based MR oximetry techniques quantitatively measure pO_2_ in tumors and muscle tissue. Recently, a novel nitroimidazole-based T1 contrast agent, gadolinium tetraazacyclododecanetetraacetic acid monoamide conjugate of 2-nitroimidazole (GdDO3NI, MW = 839 g/mol), was used to measure hypoxia, non-invasively by MRI, *in vitro* using 9L glioma cells (44) and *in vivo* in a rat prostate cancer model (45) and MRI contrast enhancement showed correlation with pimonidazole staining. The relaxivity values of GdDO3NI were reported as r_1_=4.75±0.04 s^−1^mM^−1^ and r_2_=7.52±0.07 s^−1^mM^−1^ at 37 °C and 7 T(46).

The objective of the present study is to utilize GdDO3NI enhanced MRI, for high resolution visualization of hypoxia in the rodent brain post TBI and to validate MRI data with pimonidazole based IHC. Here, we used the controlled cortical impact (CCI) injury model (47) to recapitulate elements of a focal TBI including focal lesion, axonal injury, BBB disruption, and necrosis (48,49).

## 2. Materials and Methods

### 2.1. Materials

ProHance (Gadoteridol; Bracco Diagnostics Inc., Monroe Township, NJ, USA), was used as control contrast agent. GdDO3NI, was synthesized as described previously(44). Briefly, DOTA (1,4,7,10-tetraazacyclododecane-1,4,7,10-tetraacetic acid) was selected as Gd chelator along with conjugation to 2-nitroimidazole for hypoxia targeting. Pimonidazole and FITC linked mouse anti-pimonidazole MAb (Hypoxyprobe Inc, Burlington, MA, USA) was used as the gold standard hypoxia indicator in tissue through IHC staining.

### 2.2. Animal preparation

All animal studies were approved by Arizona State University’s Institute of Animal Care and Use Committee (IACUC) and were performed in accordance with the relevant guidelines. Two cohorts of 5 animals each were used in this study. All the procedures on animals were performed under isoflurane (Baxter International Inc, Deerfield, IL). TBI was simulated using the CCI model of TBI (47). Briefly, adult male C57Bl/6 mice (9-11 weeks old) placed in stereotaxic frame under isoflurane anesthesia (3% induction, 1.5% maintenance). The frontoparietal cortex was exposed via 3 mm craniotomy and the impact tip was centered at −1.5 mm bregma and 1.5 mm lateral from midline. The impactor tip diameter was 2 mm, the impact velocity was 6.0 m/s and the depth of cortical deformation was 2 mm with 100 ms impact duration (Impact ONE; Leica Microsystems). After the impact, the skin was sutured, and the animal was catheterized for tail vein injection. Once the catheter was connected, the animal was placed onto temperature controllable MR bed and kept at 37 °C and under anesthesia (isoflurane at 1.5%).

### 2.3. Magnetic resonance imaging

MRI studies were carried out on a 7 T Bruker system with a surface coil. Mice were placed into the magnet right after the injury and pre-injection T_2_ and T_1_ weighted scans were acquired in a one hour window after injury and before injection of contrast agents. A cocktail of 60 mg/kg pimonidazole (hypoxia marker) and 0.3 mmol/kg of either MRI agents (gadoteridol or GdDO3NI) in a 100 μL volume was injected via intravenous injection at 1 hour post-injury and follow-up imaging began right after the injection. Multi gradient echo (MGE) sequence with TR= 80 ms, TE= 3 ms, flip angle= 35°, image acquisition time=5min 27s, and FOV (2 cm X 2 cm X 2 cm) were used to monitor the contrast agent uptake in the site of injury over a period of three hours with 10 min intervals after injury. Image data (k-space) were acquired with grid size of 128*64*64 and zero-filled to 128*128*128 for further analysis.

### 2.4. Data processing

Acquired data was processed using custom-written scripts in MATLAB. Regions of interests (ROI) for the muscle, injury, and the contralateral brain were manually delineating on the T_2_ maps, for each animal. These were registered and applied to the time course T_1_-wt images as masks to extract data for further analysis. From these data, the injury volume, normalized differential enhancement (NDE) and the contrast agent retention fraction, were calculated for each animal NDE is defined as the difference in percentage enhancement between the injury region and the contralateral brain and divided by the maximum value of the muscle enhancement for each data set. NDE is used to reduce the influence of normal muscle and brain contrast retention, ensuring the calculated enhancement is due to injury-induced hypoxia. The contrast agent retention fraction is calculated as the fraction of pixels in the injury region with NDE values greater than the mean NDE + standard deviation of the conventional contrast agent cohort. The contrast agent retention fraction is used to discern the additional contrast retention of GdDO3NI when compared to gadoteridol. The pre-injection and 3-hour post-injection scans were used to create 3D percentage contrast enhancement maps for each animal using the 3D Slicer software (**http://www.slicer.org**). A percentage enhancement range between 10% and 100% were used to display the 3D videos of hypoxic regions (Supplementary data) to show the spatial extent and degree of contrast retention.

### 2.5. Statistics

All results are reported as means ± standard deviations (SD). Statistical comparisons were made between time-course gadoteridol and GdDO3NI data using analysis of variance (ANOVA). In pairwise statistical evaluations, the Tukey (α= 0.05, 95% confidence intervals) test was used between specific means.

### 2.6. Tissue Collection

The excision and fixing of the brain tissue was performed as reported previously (50). Immediately after imaging, animals were deeply anesthetized with lethal dose of sodium pentobarbital solution. Once a tail pinch produced no reflex movement, animals were transcardially perfused with cold phosphate-buffered saline (PBS), followed by 4 % (w/v) buffered paraformaldehyde solution. Whole brain tissue was harvested and fixed overnight in 4 % (w/v) buffered paraformaldehyde solution. The following day, brains were immersed in 30 % (w/v) sucrose solution in 1X PBS for cryoprotection until the tissue was fully infiltrated. Samples were embedded in optimal cutting temperature medium and frozen on dry ice. Samples were stored at −80°C until sectioned coronally at a 20 μm thickness with a cryostat (CryoStar™ NX70; Thermo Fisher Scientific) and collected onto positively charged microscope slides. Slides were retained in −80oC refrigerator for further staining and analyses.

### 2.7. Histology and immunohistochemistry

Pimonidazole staining was performed using FITC-conjugated anti-pimonidazole antibody. The slides, which were kept in −80 °C were moved to the −20 °C freezer for 20-30 minutes. Then, they were moved to a 4 °C refrigerator for 15-20 minutes before being moved to room temperature PBS in a glass slide staining rack to avoid the temperature shock. The slides were moved to a tray and covered with the blocking buffer for 1 h. The slides were then washed using the waterfall technique with 1X PBS three times and were placed in the glass station filed with 1x PBS for 5 minutes and washed again. Following that, slides were placed in a tray and moved to 4 °C refrigerator and covered with 100 μL of the FITC conjugated anti pimonidazole antibody and incubation buffer mixture for overnight incubation.

Sections were then washed with 1 mL of 1X PBS for three times in a dimmed light room and incubated with the nuclear stain DAPI (4’,6-diamidino-2-phenylindole)to help in identification of tissue in final visualization. Microscope images were acquired using a Leica microscope with 355-425 nm excitation / LP 470 nm emission filter for DAPI and 450-490 nm excitation / 500-550 nm emission filter for FITC.

## 3. Results

To verify the consistency of the TBI modeling, we calculated the injury volume for each cohort using MR images. T2 wt MRI allowed for delineation of injury regions and quantification of injury volumes in the gadoteridol and GdDO3NI injected cohorts at 1 hr post-injury. The mean injury volumes for the gadoteridol and GdDO3NI cohorts were 4.82±0.50 mm^3^ and 5.23±1.13 mm^3^, respectively (p>0.05). Fig. 1 shows the dynamic percentage enhancement maps of a representative animal from the gadoteridol (top row) and GdDO3NI injected animal over the course of three hours post-injection. The region of interest (ROI) defining the injury region is indicated using a white arrow on T_2_ weighted and T_1_ weighted images. The color map overlay represents the percentage changes in pixel intensities with respect to the pre-injection value. Gadoteridol (Fig 1, top row) shows almost complete clearance from the injury region while GdDO3NI (bottom row) shows significant accumulation at 3 hr post injection. Fig. 2a shows the mean time-course NDE of the two cohorts over three hours post-injection. Statistical analysis showed significant differences (p< 0.05) in NDE of the injury region after 150 min between gadoteridol and GdDO3NI cohorts. The distribution of pooled NDE values over all animals (Fig. 2b) further underline the differences between the two agents. The ROI analysis for both contrast agents revealed changes in the signal intensity in injury site and muscle but almost no changes in the healthy contralateral region over the three hours (Supplementary figure 1). The intensity enhancement values in muscle showed rapid changes compared to the injury region that was slow and prolonged. Comparing the two cohorts (n=5) shows statistically significant retention (p < 0.05) of GdDO3NI in the injured region with significantly lower retention of gadoteridol at 150 minutes post-injection or later with the contrast agent retention factor values of 63.95±27.43 % and 20.68±7.43 %, respectively, at 3 hr post injection (4 hr post injury). In order to visualize the hypoxic regions in 3D, the maps of hypoxia and videos were generated. Fig. 3 (a, b) illustrate the 3D visualization of percentage enhancement at 3 hr post-injection for a representative animal from each cohort which shows the difference between the enhancement of each contrast agent can be seen, and the volumetric extent of the contrast agent retention. Corresponding videos (Supplementary data) allow for complete visualization of the injury region within the brain. Table 1 summarizes the injury volumes, NDE and the contrast agent retention factor for each animal as well as the means and standard deviations. We performed pimonidazole-based immunohistochemical staining of the brain sections for each cohort to study the presence of hypoxia and sections from corresponding representative animals in Fig. 2 and Fig. 3 are shown in Fig. 4. The DAPI staining is presented in blue and pimonidazole staining in green. The results from both cohorts confirmed the presence of pimonidazole in the injury region.

**Table 1.** Injury volume, normalized differential enhancement and contrast agent retention fraction for each animal in the gadoteridol (control) and GdDO3NI cohorts.

| Animal number | Injury volume (mm<sup>3</sup>) | Normalized Differential Enhancement (NDE) | Contrast agent retention fraction (%) |
| --- | --- | --- | --- |
| **Gadoteridol** |  |  |  |
| # 1 | 4.68 | -0.26 | 15.17 |
| # 2 | 5.27 | -0.04 | 13.41 |
| # 3 | 4.58 | 0.16 | 32.37 |
| # 4 | 4.20 | 0.10 | 21.71 |
| # 5 | 5.40 | -0.09 | 20.74 |
| **Mean** | **4.82±0.50** | **-0.03±0.17** | **20.68±7.43** |
| **GdDO3NI** |  |  |  |
| # 1 | 6.81 | 0.89 | 51.25 |
| # 2 | 5.66 | 0.41 | 45.32 |
| # 3 | 3.77 | 0.18 | 93.67 |
| # 4 | 4.69 | 0.18 | 36.32 |
| # 5 | 5.22 | 0.13 | 93.20 |
| **Mean** | **5.23±1.13** | **0.36±0.32\*** | **63.95±27.43\*** |

**Fig 1.**
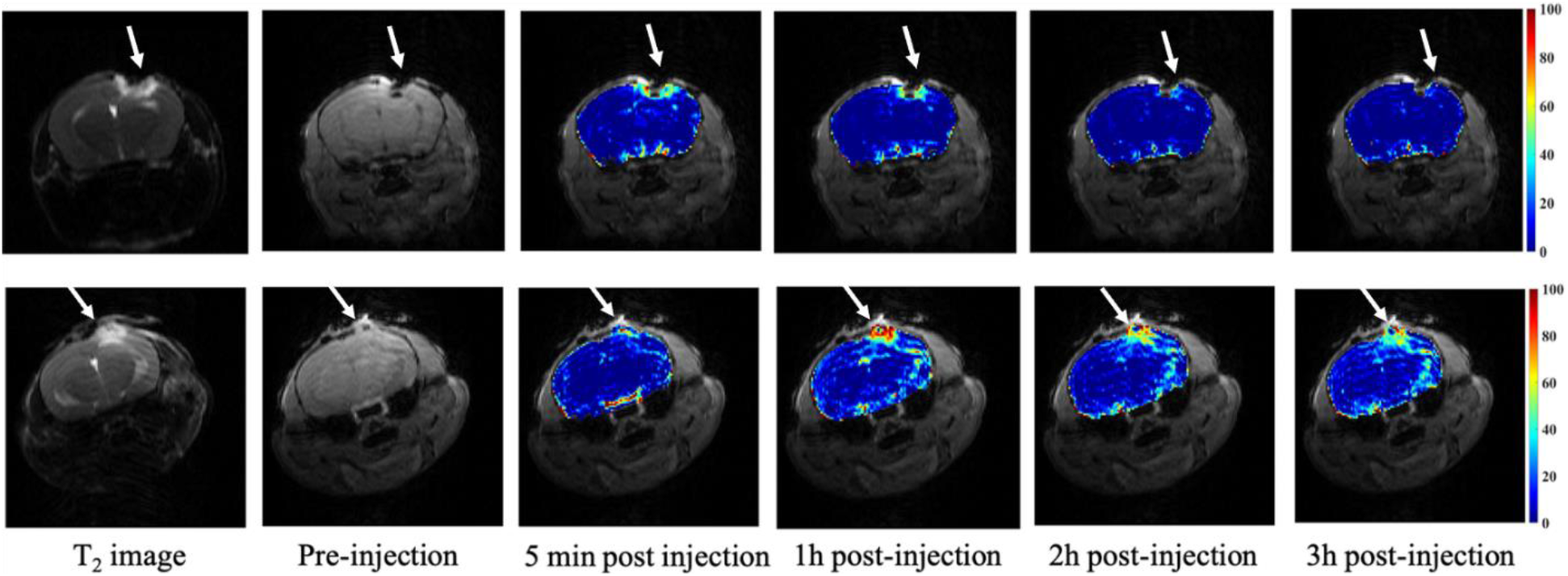
MRI T2 weighted scout images and signal enhancement (%) overlay on T1 weighted images of mice brains pre- and post-injection of conventional Gd agent gadoteridol (top row), and GdDO3NI (bottom row) over 3 hrs after injection. White arrow shows location of injury.

**Fig 2.**
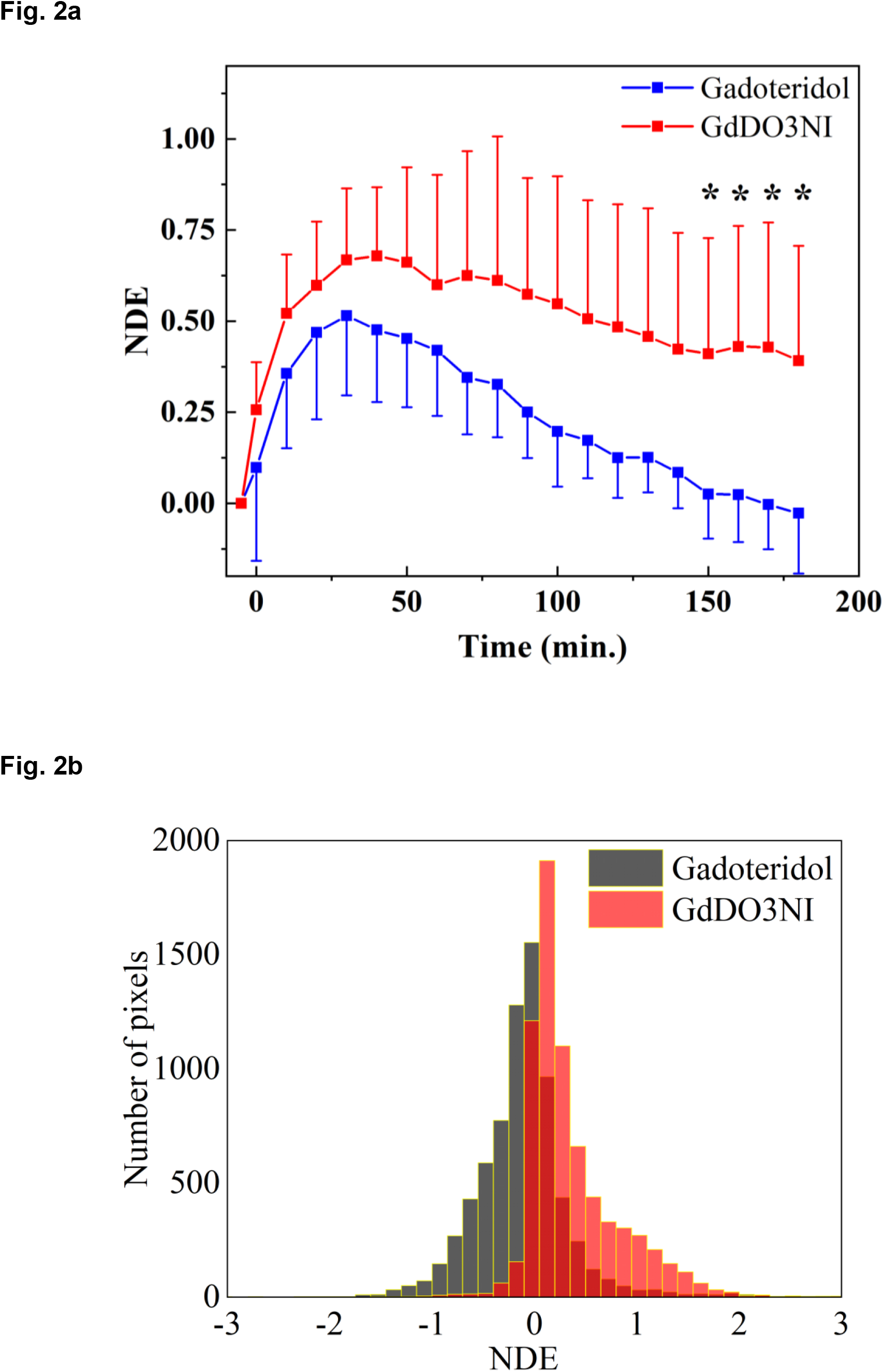
Time course mean normalized differential enhancement (NDE) results for gadoteridol and GdDO3NI injected cohort during 3 hrs after injection. NDE is calculated as the difference between the % enhancements of the injury region and a contralateral brain region of interest (ROI) normalized to the peak % enhancement for a muscle ROI.

**Fig 3.**
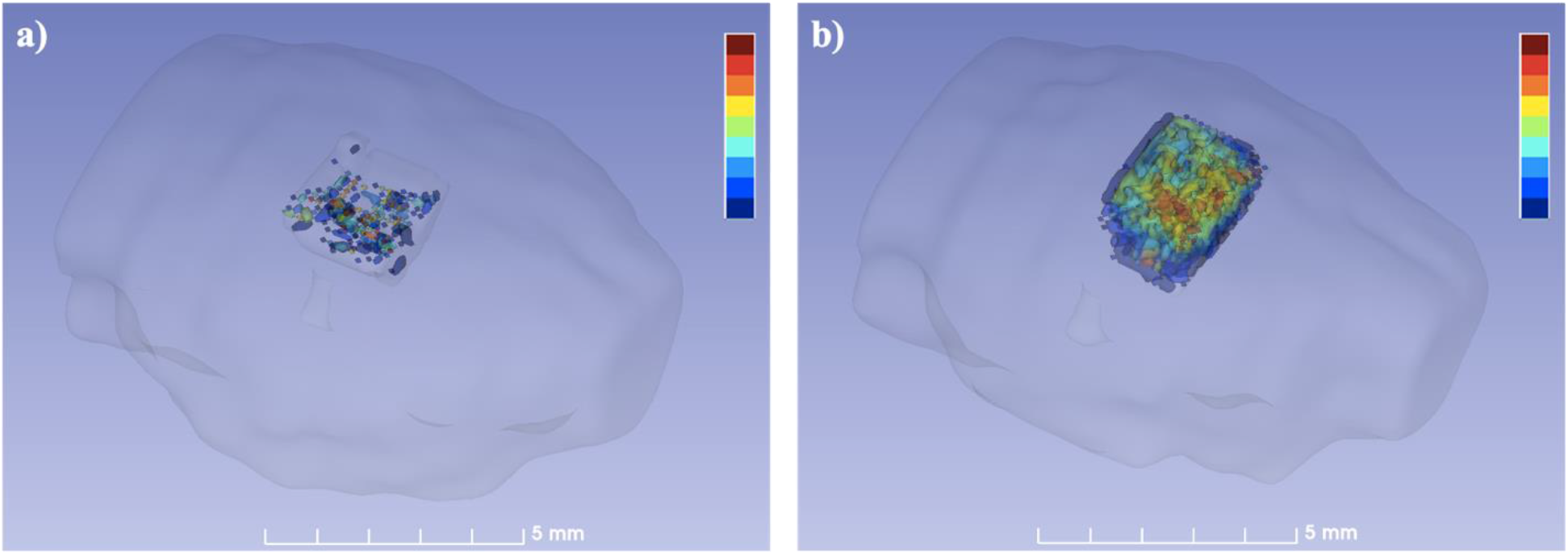
3D rendering of hypoxic regions in brains of representative (a) gadoteridol and (b) GdDO3NI injected animal. Color scale represents percentage enhancement at 3 hrs post injection of respective contrast agent. 3D rendering shows both the spatial extent as well as the severity of hypoxia in the GdDO3NI cohort and the degree of residual agent retention in the gadoteridol cohort. Color scale= 10% −100%, scale bar = 5 mm.

**Fig 4.**
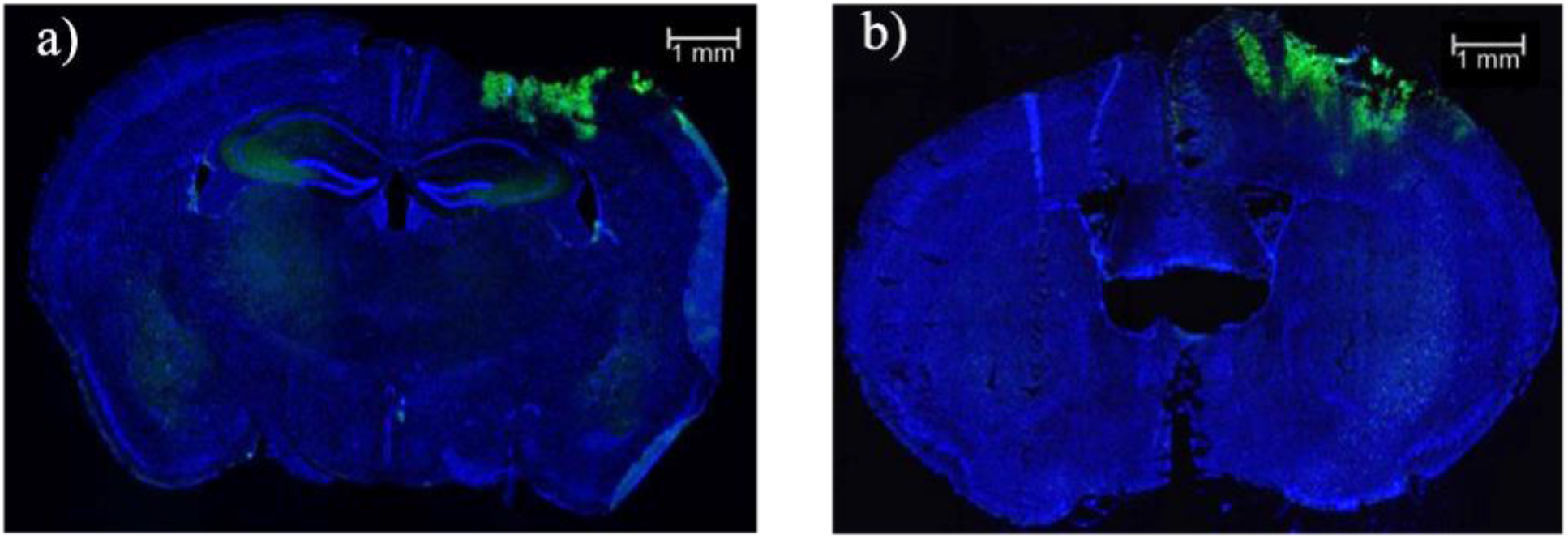
Immunohistochemical staining for hypoxia in injured animals (corresponding to fig 1), DAPI is visualized in blue and pimonidazole staining is visualized in green in representative animals injected with (a) gadoteridol or (b) GdDO3NI. Both cohorts display presence of hypoxia post injury. Scale bar = 1 mm.

## 4. Discussion and Conclusions

The mean volume of injury for gadoteridol injected cohort and the GdDO3NI injected cohort showed no statistically significant difference between the cohorts, indicating the consistency of the lesion volume and TBI modeling across the animals. Our results showed the contrast agent retention factor was significantly different in the two cohorts. Under normal physiological conditions, small molecular non-targeting Gd contrast agents such as gadoteridol, once introduced to the bloodstream, do not extravasate into brain tissue due to the tightly regulated BBB (unlike the muscle tissue). In pathologies that disrupt the BBB (e.g TBI or brain tumors), dynamic contrast enhanced MRI shows accumulation and clearance of the contrast agent from the affected region enabling the estimation of BBB leakiness (51,52). GdDO3NI and gadoteridol had similar, slower uptake pharmacokinetics in the brain injury region compared to muscle in each case. This observation can be attributed to the complex dynamic of the injured brain and blood flow disruption due to BBB dysfunction in the injury region compared to the intact muscle (53). In contrast, the clearance pharmacokinetics of the two agents (GdDO3NI and gadoteridol) were markedly distinct in the injured brain while they were similar in the muscle region. In absence of hypoxia in the muscle, GdDO3NI behaves like any other small molecular MRI contrast agent, including gadoteridol) with uptake/clearance kinetics that are predominantly flow-limited (54). However, in the hypoxic injured region, the two agents behave differently, with GdDO3NI binding to proteins, via the 2-nitroimidazole moiety of the agent, showing significantly longer retention times compared to gadoteridol. This mechanism has been previously shown in prostate tumors as well where GdDO3NI was retained in hypoxic regions compared to a non-targeting Gd agent (45). A qualitative comparison between the MRI and IHC images shows a good agreement between the location of T1 hyper-intensity regions within in the injury site in the GdDO3NI cohort and pimonidazole binding. These observations confirm our hypothesis that GdDO3NI localizes in hypoxic regions in TBI.

The contrast agent retention factor reflects the amount of contrast agent in the injury region at the 3 hr post injection. While we see significant differences in the retention of gadoteridol compared to GdDO3NI, the former does not appear to be completely eliminated possibly due to the irreversible retention (51). The dynamic nature of the BBB dysfunction can influence the amount of exogenous agents that are able to wash in or wash out, even over a matter of hours post injury, due to dynamic reduction in BBB leakiness (55). Our results showed the GdDO3NI cohort had significantly higher retention values compared to gadoteridol that can only be attributed to binding in the hypoxic regions. In case of the GdDO3NI the retention fraction is related to the hypoxic fraction but, unlike tumors, it may overestimate the hypoxic fraction due to some degree of irreversible retention arising from acute reduction in BBB leakiness over 3 hrs since the contrast agent was injected (as seen for the control agent, gadoteridol). The IHC results from both cohorts confirmed the presence of hypoxia post TBI; however, only animals in the GdDO3NI cohort were able to represent that in MRI contrast enhancement studies.

In summary, the results demonstrate that contrast enhanced MRI using GdDO3NI allows visualization of hypoxic regions in the brain following TBI. The MR results were validated by the gold standard method of IHC staining for pimonidazole. Non-invasive imaging of hypoxia in TBI could allow for injury prognosis as well as personalized treatment targeted towards alleviation of hypoxia.

## ACKNOWLEDGMENTS

This study was supported in part by a National Science Foundation CAREER Award#1351992 (V. D. K), a “Rising Stars in Engineering” seed grant from the Fulton Schools of Engineering, ASU (V.D.K., S.E.S.), a FLINN Foundation grant (S.E.S., V.D.K.), NIH 1DP2HD084067 (S.E.S.) and Arizona State University Graduate College Completion Fellowship (B.M. and V.N.B.). Imaging studies were performed at the Barrow-ASU Center for Preclinical Imaging, a part of the Biosciences Core Facilities at Arizona State University.

