## Supplemental data for "GdDO3NI allows imaging of hypoxia after brain injury"

### Table of content

Intensity enhancement images of the Gadoteridol cohort……………………….....…..... ii

Intensity enhancement images of the GdDO3NI cohort………………………………. iv

Video of 3D representation of contrast agent retention……………………………….... vi

**Figure S1.** Signal intensity enhancement data in the injured brain, contralateral brain, and muscle regions (%) for the individual animals in the Gadoteridol cohort. Gadoteridol, the conventional, nontargeting contrast agent served as control in this study. The ROI analysis was a voxel-wise comparison and was performed in MATLAB by measuring the percentage enhancement of signal intensity in $T_1$ weighted images over three hours of the experiment compared to the pre-injection image.

Animal #1

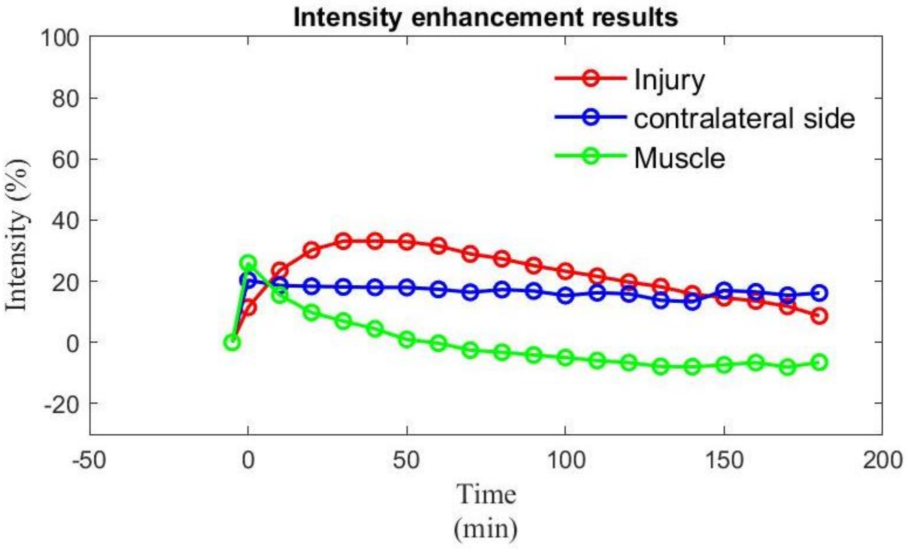


Animal #2

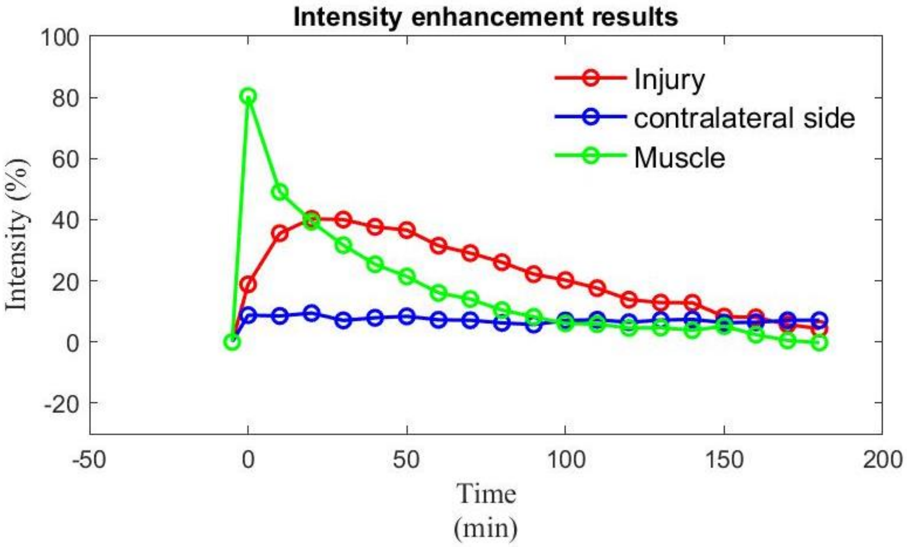

Animal #3

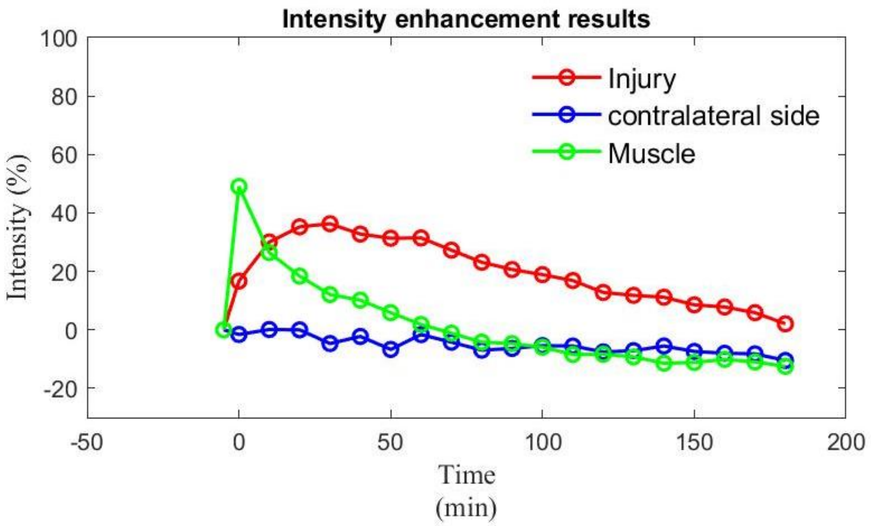


Animal #4

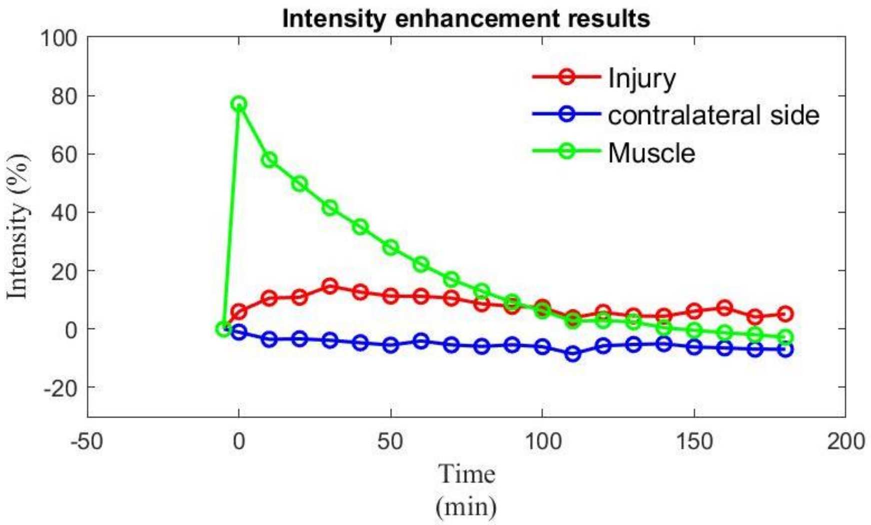


Animal #5

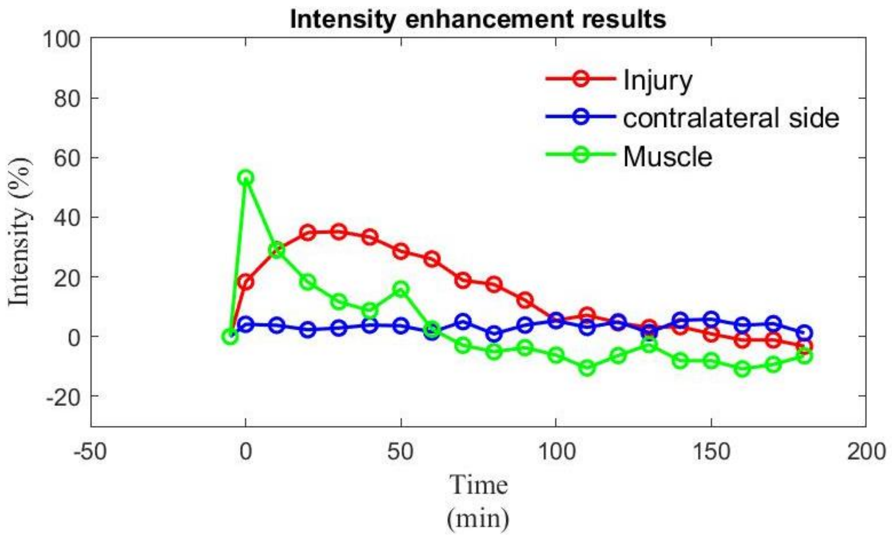

**Figure S2.** Signal intensity enhancement data in the injured brain, contralateral brain, and muscle regions (%) for the individual animals in the GdDO3NI cohort. The ROI analysis was a voxel-wise comparison and was performed in MATLAB by measuring the percentage enhancement of signal intensity in $T_1$ weighted images over three hours of the experiment compared to the pre-injection image.

Animal #1

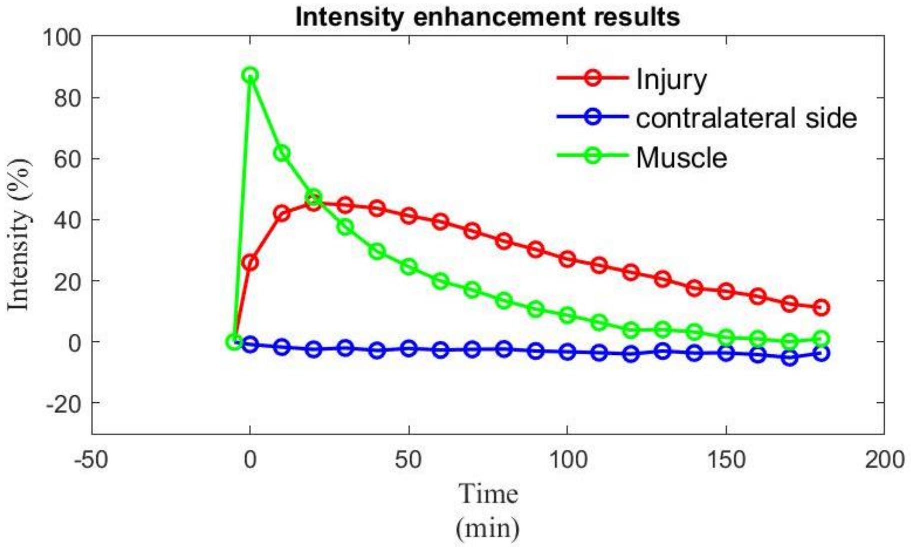


Animal #2

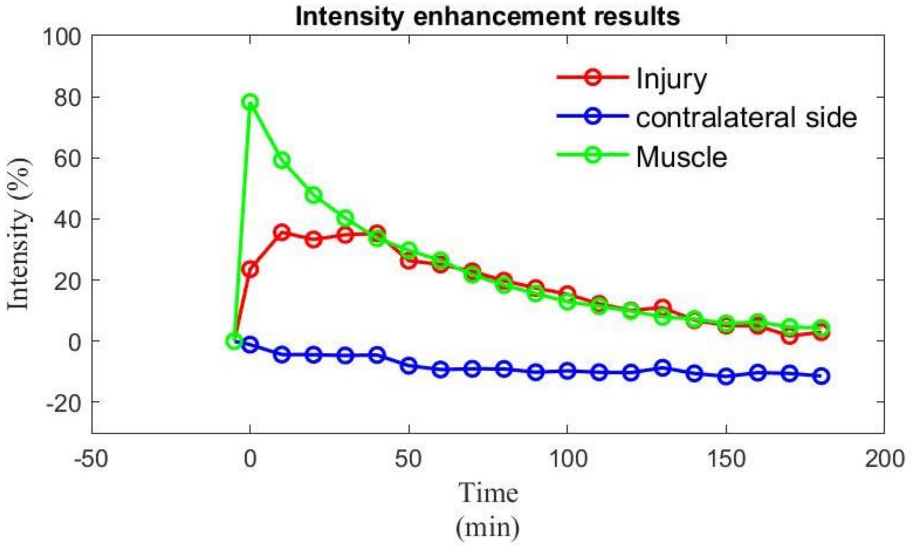

Animal #3

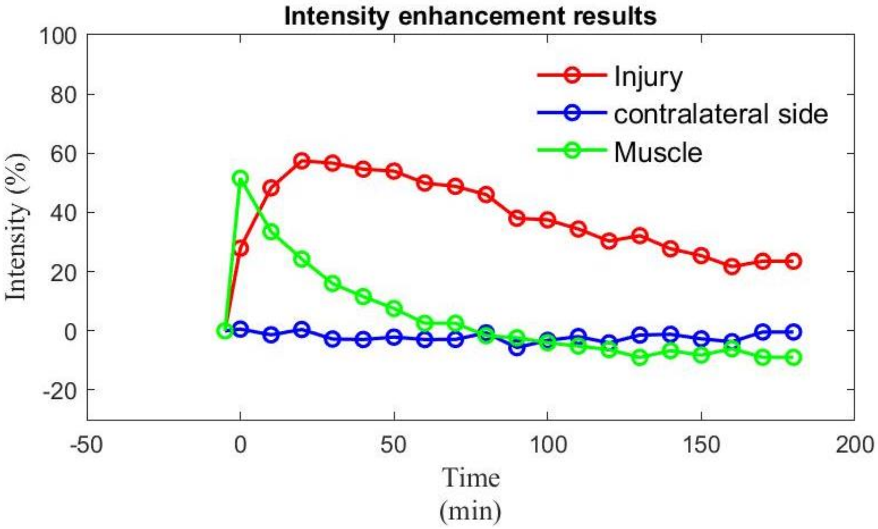


Animal #4

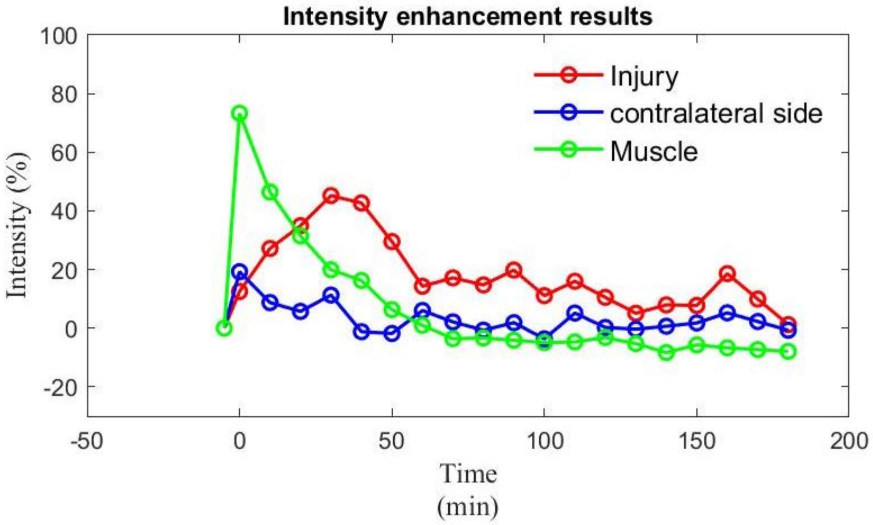


Animal #5

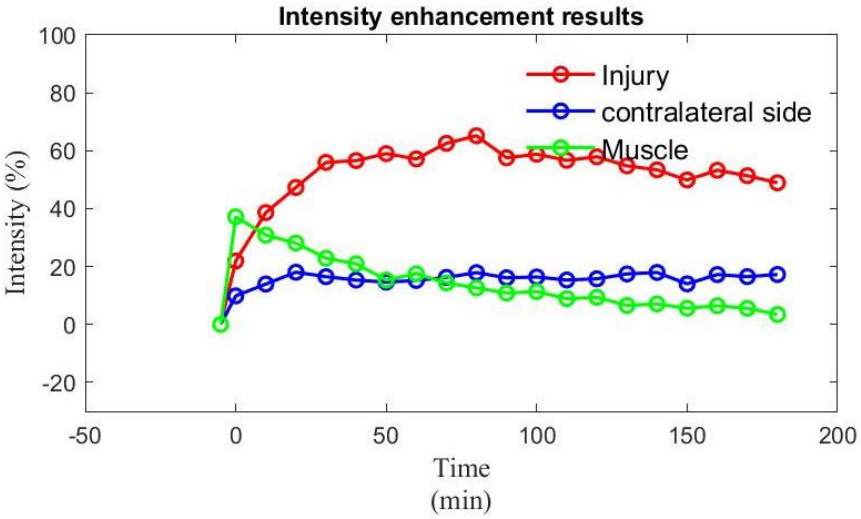

**Video S1.** The brain and injury region models were created from the manually-created ROIs. Contrast enhancement percentages between 10% and 100% were used to create models of hypoxic regions, segmented by the intensity of the contrast enhancement. The segments were colored following a JET color scheme where blues represent lesser enhancement and reds represent greater enhancement. Each segment contains a 10% range of percent enhancements, creating nine segmentations between 10% and 100%. These models are presented together to form a cohesive view of hypoxia within traumatic brain injuries. 3D rendered and quantified contrast percentage enhancement, comparing contrast agent retention a) Gadoteridol and b) GdDO3NI.

a) Gadoteridol

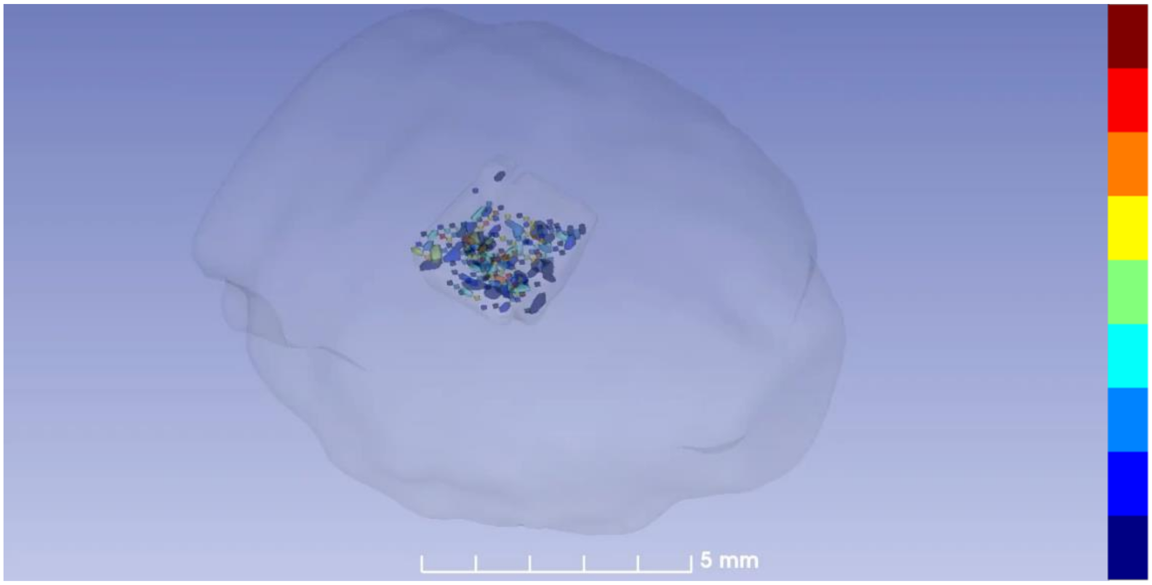


b) GdDO3NI

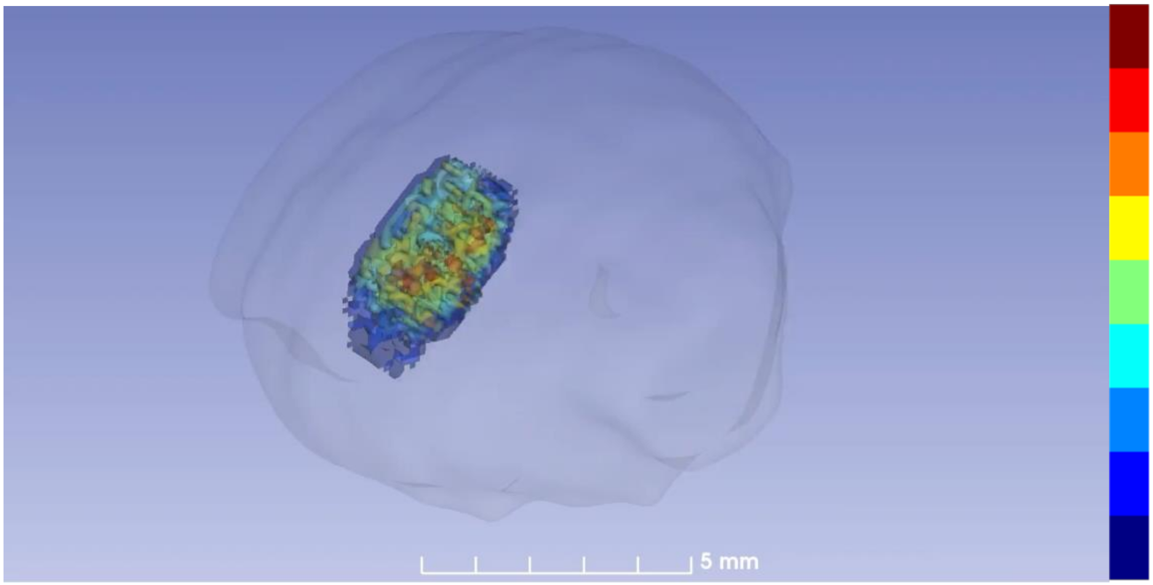
